# Microbial fingerprints reveal interaction between museum objects, curators and visitors

**DOI:** 10.1101/2023.01.04.522779

**Authors:** Lukas M Simon, Cecilia Flocco, David Henke, Luise Rauer, Christian Müller, Johannes Vogel, Christiane Quassier, Jörg Overmann, Stefan Simon

## Abstract

Microbiomes populate the border between humans and their environment. Whether the microbiome can be leveraged to gain information on human interaction with museum objects is unclear. To answer this question, museum objects varying in material and size from two museums, the Museum für Naturkunde and the Pergamonmuseum in Berlin, Germany, which forms part of UNESCO World Heritage since 1999, were defined. In total 126 samples of natural and cultural heritage objects were taken with sterile nylon flocked swabs and subsequently subjected to 16S rRNA amplicon sequencing. By comparing the microbial composition of touched and untouched mollusc and fossil natural heritage objects we derived a robust microbial touch signature characterized by increased abundance of microbes known to be present in human skin. Application of this touch signature to cultural heritage objects from the Pergamonmuseum revealed areas of differential exposure to human contact on the Ishtar gate and Sam’al gate lions. Moreover, we were able to distinguish museum objects and personal office items touched by two different individuals with high sensitivity. Our results demonstrate that the microbial composition of museum objects gives insight into the degree of exposure to human contact, which is an important parameter for conservation and heritage science, and possibly provenance research.

## Introduction

In light of the global debate on ownership in collections, both of cultural and natural heritage, the significance of provenance research has dramatically increased over the past years. Furthermore, the authenticity of art objects and the ability to reliably distinguish fakes from originals, is equally important. More and more, in the light of the post-colonial discourse, museum visitors and the wider public take a growing interest in these questions as well. Apart from deterioration processes, the interaction between object and environment is reflected in the composition of the associated microbial communities. A better knowledge of this relation will significantly advance the research of provenance in natural history and cultural collections, open avenues to new knowledge on population dynamics, co-evolution and symbiosis, and geographical and climatic conditions. Although different in specific aspects, the main challenges of cultural and natural heritage are the same.

Microbiomes, which refer to the community of microorganisms that live on or within an object, act as indicator for interactions - between organisms (Meadow et al. 2013; Moeller et al. 2016), organisms and objects (Fierer et al. 2010), objects and their environment (Senghor et al. 2018). Microbiome analysis meanwhile is a well-established approach in biology and medical studies (Consortium and The Human Microbiome Project Consortium 2012). On the level of the human individual, microbiomes provide insights in the personal health status and possible causes of dysfunctions. On the community level, the microbial landscape and microbial load on public places are used as an indication for the health risk given through the identification of certain bacteria or other microbes (Afshinnekoo et al. 2015). Cities possess a consistent “core” set of non-human microbes, which reflect important features of cities and city-life (“A Global Metagenomic Map of Urban Microbiomes and Antimicrobial Resistance” 2021). With the growing potential to detect a “molecular echo” of an individual’s microbiome on touched surfaces, which could lead to substantial progress in the scientific analysis of forgery and illicit traffic cases in the field of cultural heritage (Simon and Röhrs 2018), new ethical and legal aspects need to be considered (Elhaik et al. 2021).

In the field of heritage science, microbiome analyses are used to identify and assess deterioration processes in porous building materials or cellulose-based substrates, parchment, paintings (Marvasi et al. 2019). So for example halophilic microorganisms, involved in the biodeterioration of historic building materials, such as brick and paint coating, have been identified by (Adamiak et al. 2017). Specific genera such as Aeromonas present on canvas and panel paintings can be potentially responsible for deterioration and fading of paint layers (Torralba et al. 2021). Microbial biofilms are known to cause health issues and also to catalyze deterioration processes (Sterflinger and Piñar 2013).

Studies of microbiomes on smaller, mobile heritage objects are still rare. Some are e.g. dealing with health risk of handling heritage objects (Górny et al. 2016) and the microbial profile on historical books and archives ([MIKROBIB project](https://www.tu-braunschweig.de/en/philosophie/mikrobib)). In a study on smuggled marble sculpture of unknown origin, microbiome analysis was performed with the aim to reconstruct the history of the storage of the objects, to infer a possible relationship among them, and to elucidate their geographical shift (Piñar et al. 2019). In each of these cases, the microbiome being sampled is the product of ongoing or recent processes with the environment.

Here, we analyzed and assessed the microbiome on natural and cultural heritage objects in two museums, the Berlin Museum für Naturkunde (MfN) and the Pergamonmuseum (PM), part of the Stiftung Preussischer Kulturbesitz (SPK) and UNESCO World Heritage since 1999, at the crossroad of art and science, to answer the following questions:

Is there a robust microbial signature of the environment in which the objects are stored? How long does this signal perdure in the face of handling efforts, or visitor exposure, which may produce different biological signals? Are there any indications for (living) curators or the general public on the objects they interacted with? By combining data science techniques with microbial sampling approaches we obtained and interpreted the microbial profile from heritage objects to generate a reliable baseline for decoding new knowledge.

## Results

### 16S rRNA sequencing captures microbial profile of museum objects

To assess the feasibility of detecting interactions between museum objects and their environment, minimally invasive microbial analysis of museum objects from the Pergamonmuseum (PM) and Museum für Naturkunde (MfN) in Berlin was conducted through swabbing. We selected museum objects of varying size and materials, in various locations (exhibition, collection, workshop) which allowed sampling of areas with variable exposure to human contact (**Table 1**).

**Table 1.**
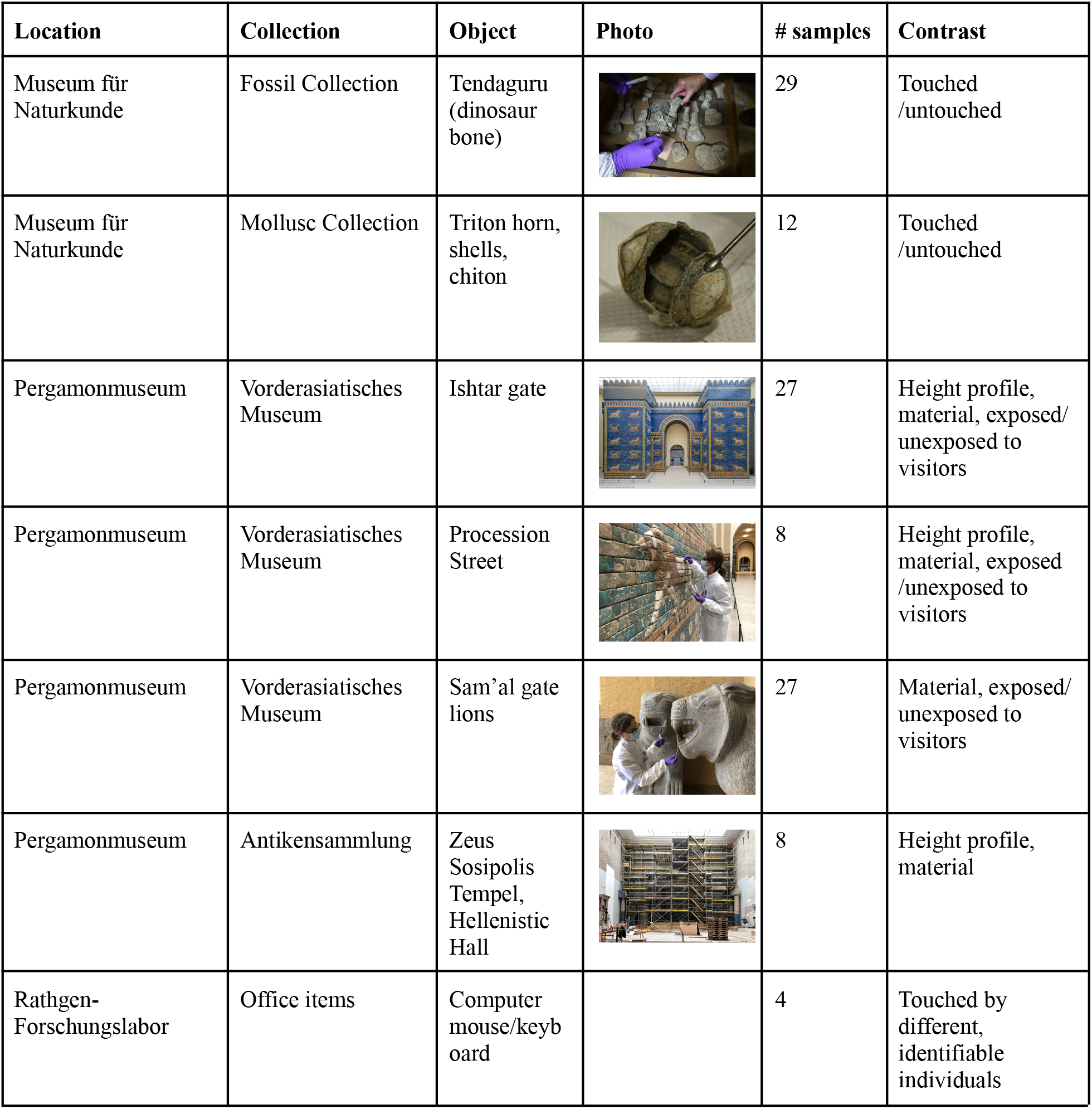
Overview of sampled objects.

| Location | Collection | Object | Photo | # samples | Contrast |
| --- | --- | --- | --- | --- | --- |
| Museum für Naturkunde | Fossil Collection | Tendaguru (dinosaur bone) |  | 29 | Touched /untouched |
| Museum für Naturkunde | Mollusc Collection | Triton horn, shells, chiton |  | 12 | Touched /untouched |
| Pergamonmuseum | Vorderasiatisches Museum | Ishtar gate |  | 27 | Height profile, material, exposed/ unexposed to visitors |
| Pergamonmuseum | Vorderasiatisches Museum | Procession Street |  | 8 | Height profile, material, exposed /unexposed to visitors |
| Pergamonmuseum | Vorderasiatisches Museum | Sam'al gate lions |  | 27 | Material, exposed/ unexposed to visitors |
| Pergamonmuseum | Antikensammlung | Zeus Sosipolis Tempel, Hellenistic Hall |  | 8 | Height profile, material |
| Rathgen-Forschungslabor | Office items | Computer mouse/keyb oard |  | 4 | Touched by different, identifiable individuals |

Samples were collected during two sampling rounds in October 2020 and 2021, respectively. To obtain the biological information without damaging the objects in any manner, sampling was carried out by wiping DNA-free and sterile nylon flocked swabs across the surface. A total of 126 samples from unique museum objects and controls was collected.

Samples were subsequently subjected to 16S rRNA amplicon sequencing. To keep microbial discovery parameters consistent, raw sequencing data from both sampling rounds were merged and processed together (**Figure 1A, S1**). The resulting processed count tables quantify the abundance of Amplicon Sequence Variants (ASV), which represent DNA sequences detected in each sample. Each ASV feature may or may not be assigned to a phylogenetic tree with variable resolution.

**Figure 1.**
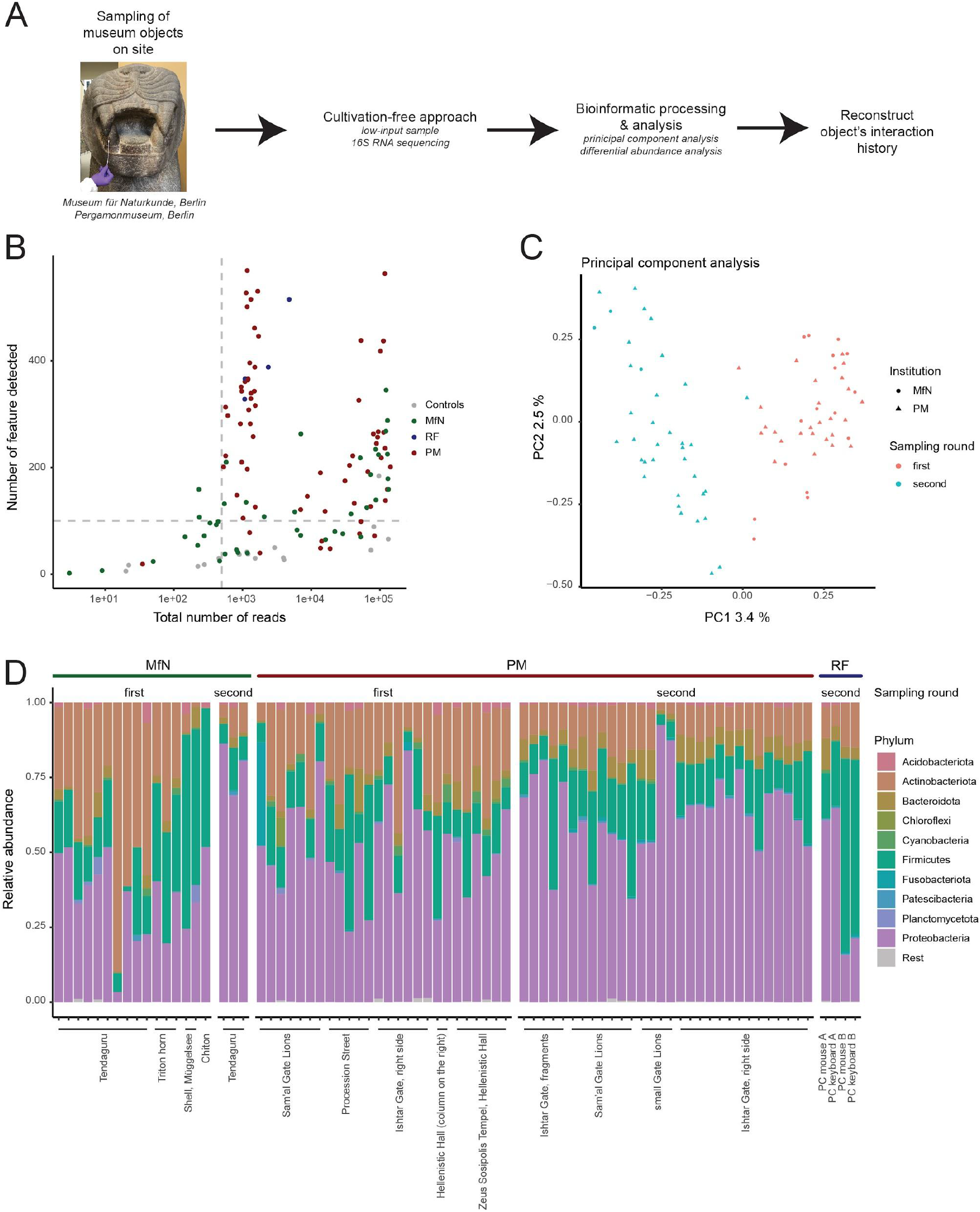
16S rRNA sequencing captures microbial profile of museum objects. A) Workflow overview. B) Scatter plot shows the total number of reads (x-axis) and number of unique ASV features detected (y-axis) across all samples and controls. Samples taken at the MfN and SPK are colored blue and red, respectively. Negative control samples are colored green. Grey dashed lines indicate quality control thresholds. C) Principal components 1 (x-axis) and 2 (y-axis) stratify samples and controls. Samples taken during the first and second sampling rounds are colored red and blue, respectively. D) Barplot shows relative abundance of microbial phyla (y-axis) across all samples passing quality control filtering (x-axis). Colors represent the ten most frequent phyla.

On average each sample contained a total of 31,446 reads with 194 unique ASVs detected. Given that many samples contained relatively little DNA, we used a set of negative controls to define quality control filtering criteria. These negative controls included samples taken of air or samples including only various library preparation reagents. While the negative control samples showed a large range of total reads, only a single negative control sample had more than 100 ASV features detected (**Figure 1B**). Therefore, we focused analysis on a total of 79 samples with more than 100 unique ASV features detected and more than 500 total reads excluding negative controls.

To explore the data, we applied unsupervised dimension reduction using principal component analysis to the 79 samples passing quality control filtering (Figure 1C). We observed that the first principal component separated samples taken during the first and the second sampling round indicating the presence of ‘batch effects’. Batch effects represent nuisance factors commonly observed in molecular data (Kim et al. 2017). Therefore, we limited most comparisons to within each sampling round. Investigating the phylogenetic makeup of the microbial features revealed that the majority of reads originated from phyla of *Proteobacteria, Firmicutes* and *Actinobacteria* (Figure 1D).

### Microbial fingerprint separates touched and untouched natural history objects

We first analyzed natural history objects from the MfN obtained during the first sampling round from the mollusc and fossil collections consisting of three dinosaur bones from Tendaguru (Lindi region, Tanzania), one shell of a triton (“Triton horn”, Ranellidae), one Green Chiton (*Chiton olivacaeus*), and one mussel shell collected in the Lake Müggelsee near Berlin (**Figure 2A**). A total of 15 samples passed quality control filtering and were included in this analysis. Some of these objects were commonly used as hands-on examples for visitors, while others were stored in the archives away from human contact. Thus, some objects were frequently touched by visitors and curators while others remained largely untouched. Therefore, these museum objects provided the opportunity to ask questions about the relationship between the object’s environment and its microbiome composition.

**Figure 2.**
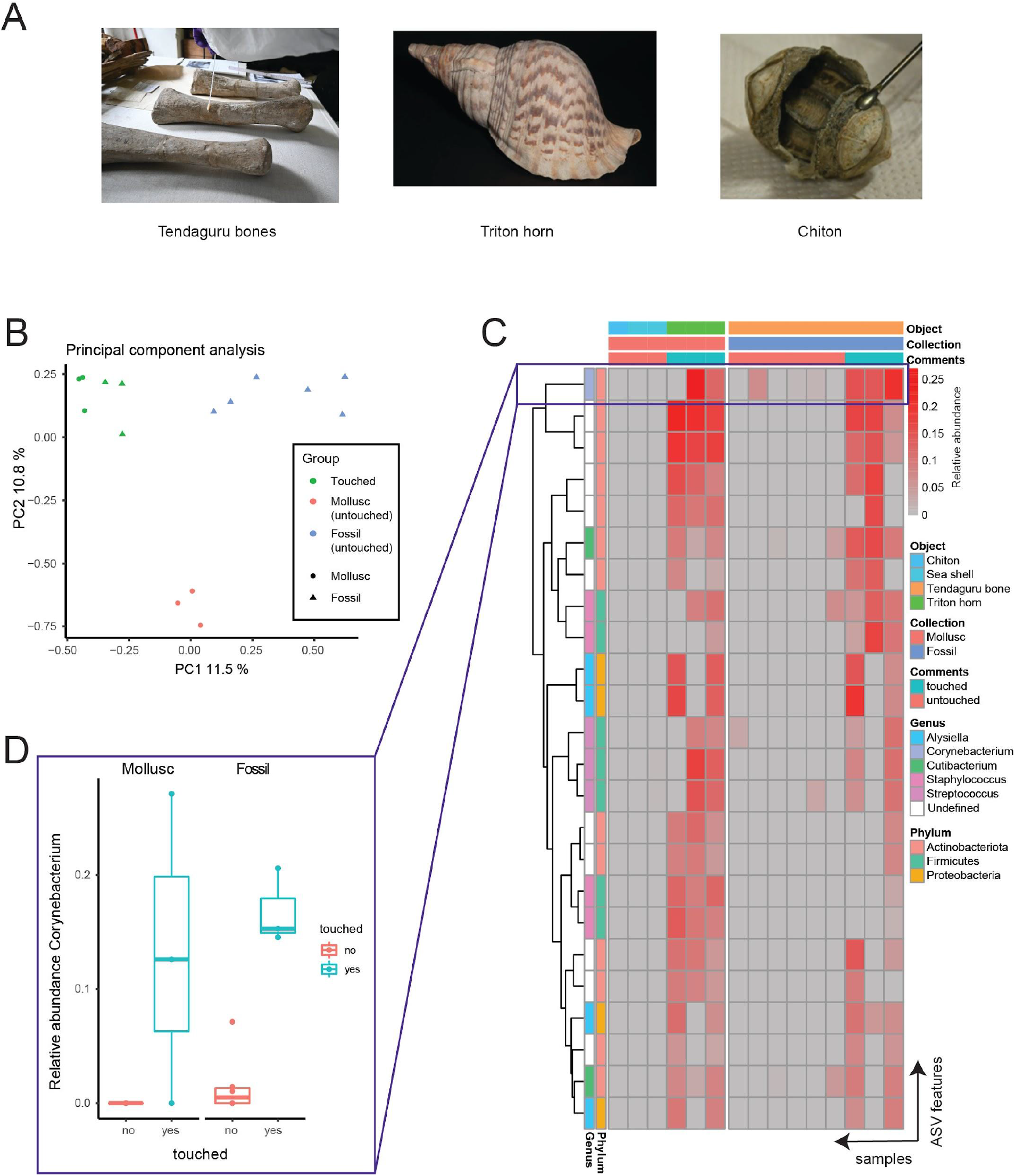
Microbial fingerprint separates touched and untouched natural history objects. A) Photos shows sampled natural science museum objects. B) Principal components one (x-axis) and two (y-axis) separate touched (green) from untouched molluscs (red) and fossils (blue). Numbers represent the proportion of variance explained by each component. Colors and shapes represent touch status and collection, respectively. C) Heatmap illustrates abundance of ASV features (rows) significantly positively associated with touch status across samples (columns). Grey and red colors represent low and high abundance values. D) Boxplot displays the abundance of an ASV feature assigned to Cornyebacterium at the genus level across touched (blue) and untouched (red) samples for both molluscs (left) and fossils (right). This genus is known to be highly abundant in human skin. Purple boxes mark data in heatmap panel C corresponding to data shown in panel D.

Unsupervised dimension reduction using principal component analysis revealed that frequently touched objects from both mollusc and fossil collections converged into a single cluster while untouched mollusc and fossil objects clustered separately (**Figure 2B**). These results suggested that the microbial profiles of untouched mollusc and fossil objects were distinct but upon touch converged onto an identical microbial profile.

Next, we set out to discover the taxonomic identities of the ASV features driving the separation between touched and untouched objects. Towards this end, we conducted differential abundance analysis comparing the abundance of each ASV feature between the touched and untouched objects. A total of 24 ASV features were significantly positively associated with touch status (Multivariate regression, FDR < 25%, **Figure 2C**). For example, an ASV feature assigned to *Cornyebacterium* at the genus level was almost exclusively detected in both touched mollusc and fossil objects compared to untouched samples (Multivariate regression, P < 0.001, **Figure 2D**).

Given that we observed a strong effect of touch on the composition of the microbiome of these museum objects, we derived a touch signature by calculating the proportion of reads assigned to any of these 24 ASV features. This touch signature robustly separated touched from untouched mollusc and fossil objects (Figure S2).

Importantly, several features in this touch signature were previously linked to the human skin microbiome (Byrd, Belkaid, and Segre 2018). In particular, *Corynebacterium, Staphylococcus*, and *Streptococcus* were among the most abundant bacterial genera of the human skin. Our taxonomic classifier was able to assign genus level information for 14 out of the 24 ASV features included in our signature. 8 of those 14 ASV features were derived from one of these three genera associated with human skin. These results further validate that the microbial touch signature we identified on the museum objects indeed represents interaction with human skin.

### Microbial fingerprint reflects exposure to human contact in antiquity objects

We next sought out to investigate the microbial profile of cultural heritage objects from the Pergamonmuseum. The Ishtar Gate was a grand entrance to the ancient city of Babylon. The gate was built during the reign of King Nebuchadnezzar II in the 6th century BCE and was decorated with glazed brick reliefs depicting dragons, bulls, and lions. Only small fragments were found during the excavation by the Deutsche Orient-Gesellschaft (German Oriental Society) from 1899 to 1917 in Babylon. Friedrich Rathgen, first Director of the Rathgen-Forschungslabor, planned and carried out an ambitious desalination campaign for the fragments, before the animal reliefs were re-assembled, supplemented with modern glazed bricks, and presented to the public when the Pergamonmuseum opened in 1930.

We first compared a total of 10 samples, taken of ceramic tiles at varying heights ranging from 1.2 to 7.6 meters (**Figure 3A**). Following principal component analysis of this subset of samples, we discovered a significant association between the first principal component and height, indicating that the microbial profile of the tiles varied with height in a continuous manner (Pearson correlation, Rho: 0.98, P: 5.8e-7) (**Figure 3B**). Of note, no significant association was observed between principal components one and two with either color (blue versus yellow) or illustration of the tiles (dragon or bull) (**Figure S3**). This is of special interest, as often blue tiles are not original, but socalled “Nachbrandziegel”, dating to the 20th century time of reconstruction of the gate.

**Figure 3.**
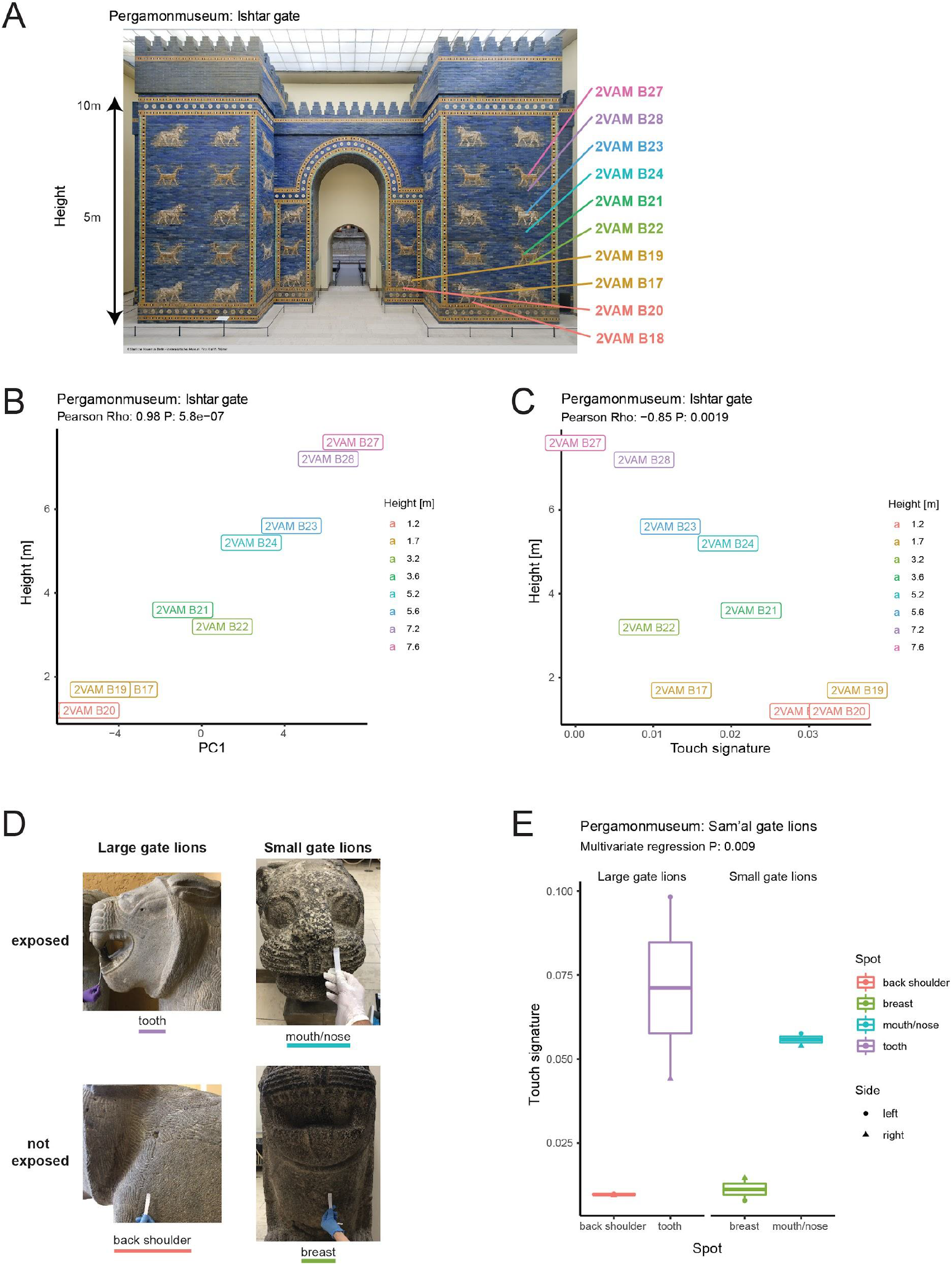
Microbial fingerprint reflects exposure to human contact in antiquity objects. A) Photo shows location of sampling locations on Ishtar gate. Photo credit: Staatliche Museen zu Berlin - Vorderasiatisches Museum, Olaf M. Teßmer. B) Principal component one (x-axis) correlates with height (y-axis). C) Touch signature (x-axis) correlates negatively with height (y-axis). For B) and C), Samples are colored by height. D) Photos depict sampling spots with varying degree of exposure to human contact for both large and small gate lions Photo credit: Rathgen-Forschungslabor, Stefan Simon. E) Boxplot illustrates differences in touch signature levels (y-axis) across various sampling areas of gate lion sculptures (x-axis). Color represents sampling areas and shape represents the side of the museum hallway where the sculpture is located.

While no single ASV feature passed significance filtering after multiple testing correction, the top 30 most strongly associated ASV features showed differential abundance with height (**Figure S4**). This observation suggests that the signal of any single ASV feature may not be strong enough to reach statistical significance given the limited sample size. However, the signal is enhanced and reaches statistical significance when many ASV features are aggregated together as captured by principal component one.

Given that lower tiles are within reach of the museum visitors we hypothesized that there is an association between height and the exposure to human contact. As a proxy for exposure to human contact we used our touch signature derived from the analysis of mollusc and fossil objects. Indeed, the touch signature significantly decreased with height (Pearson correlation, Rho: −0.85, P: 0.002) (**Figure 3C**), indicating that lower tiles had greater exposure to human contact compared to higher tiles.

Having identified an association with the touch signature and height of the Ishar gate, we asked if we could identify variable exposure to human contact on a smaller antiquity object. Therefore, we analyzed a total of four stone sculptures of the Sam’al gate lions, a pair of small and large gate lions on each side of the museum hallway. The mouth/nose and the tooth area of the small and large gate lions, respectively, are exposed and may get touched by visitors. In contrast, the breast and back shoulder areas of the small and large gate lions, respectively, are less exposed and less attractive to physical interaction with visitors. (**Figure 3D**). Indeed, the touch signature was significantly higher for areas with greater exposure to visitors across both small and large gate lions (Multivariate regression, P: 0.009) (**Figure 3E**).

### Microbial fingerprint separates objects touched by different persons

Taken together, these observations suggested that one of the major sources of variation in the microbial profiles of museum objects is the exposure to human contact. Next, we set out to test whether the microbial profile could be used to differentiate objects touched by two different persons. Towards this aim, we sampled personal office items including computer mouse and keyboard frequently and solely touched by two different subjects, called subject A and B. These office items served as positive controls because we expect to find a high load of personal microbes on them which increases the sensitivity for distinguishing two subjects. In addition, each subject touched the surface of glazed ceramic tiles from the Ishtar gate with lower levels of exposure to visitors and the back shoulder of the Sam’al gate lions made out of stone immediately prior to sampling for a period of 60 seconds. As an additional control, one tile from the Ishtar gate with high levels of exposure was touched by subject B prior to sampling.

Independent principal component analysis of these 11 samples revealed that the first principal component explained 16% of variance and separated objects based on whether it was touched by subject A or subject B (**Figure 4AB**). The tile with exposure to visitors and touched by subject B, clustered with objects touched by subject B, indicating a loss of the visitor fingerprint. The second principal component explained only 14% of variance and separated the office items from the museum objects (**Figure 4C**). Of note the office items consist of a cast polymer while the Ishtar gate tiles are made of glazed ceramic and the Sam’al gate lions are made of stone. Moreover, the museum objects are housed in the exhibition hall of the Pergamonmuseum while the office items come from the Rathgen Research Laboratory.

**Figure 4.**
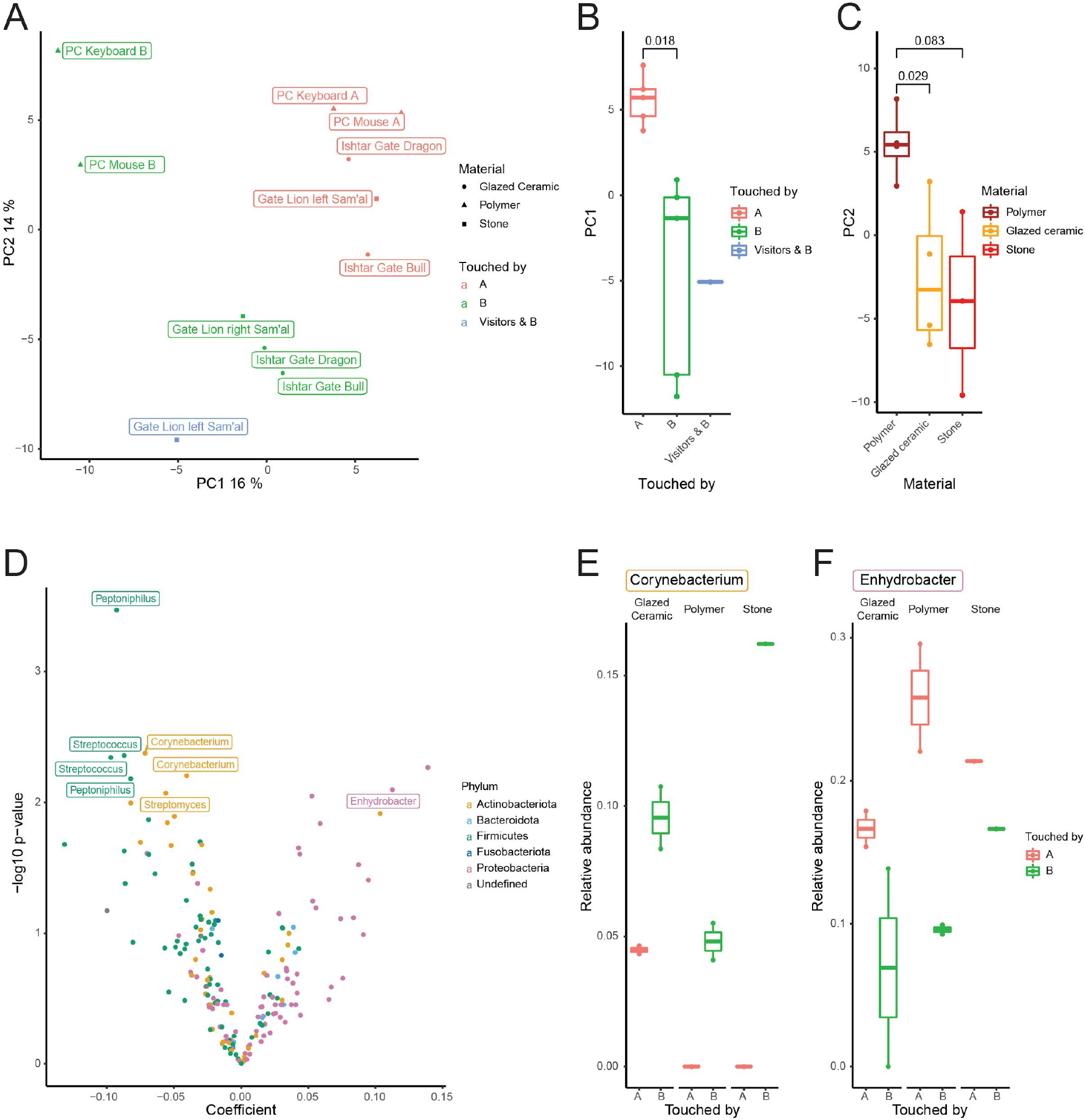
Microbial fingerprint separates objects touched by different persons. A) Principal components 1 (x-axis) and 2 (y-axis) separate objects touched by different subjects. Colors represent which subject touched the object. Shape represents the material of the object. B) Boxplot shows the principal component one (y-axis) stratified by which subject touched the object (x-axis). C) Boxplot shows principal component two (y-axis) stratified by the material of the object (x-axis). For B) and C), p-values are derived from two-sided T-tests. D) Volcano plot shows the coefficient (x-axis) derived from the multivariate regression model and the -log10 transformed p-value (y-axis) for ASV features. Colors represent Phylum. Available Genus information for top 10 ASV features is displayed. Positive coefficient values indicate higher abundance of ASV feature in subject A compared to B. E) and F) Boxplots depict differences in abundance of exemplary ASV features. Abundance of ASV features assigned to Cornyebacterium and Enhydrobacter at the Genus level is greater and lower in subject B compared to subject A, respectively.

Next, we performed differential abundance analysis. We compared the abundance of ASV features between subject A and B while accounting for the material of each object using multivariate regression. Indeed, several ASV features were identified with significantly altered abundance in subject A compared to subject B (Multivariate regression, FDR < 25%) (**Figure 4D**). For example, two ASV features assigned to *Cornyebacterium* and *Enhydrobacter* at the genus level showed lower and higher abundance in subject A compared to B, respectively (**Figure 4EF**). Of note, the differential abundance was robustly detected across multiple materials for the objects from the Ishtar gate, gate lion sculptures, and office items.

## Discussion

Taken together our results demonstrate that human touch or exposure to human contact are major contributors to the variance in microbial composition observed on the surfaces of museum objects. Moreover, human touch impacts the microbial composition of objects to a degree that enables distinction between objects touched by two different individuals. Our results suggest that other variables including material, paint and location explain relatively smaller proportions of variance.

Given the requirement of minimally invasive sampling methodology, we applied dry nylon swabs to sample the objects. Thus, the amount of DNA obtained from each sample was at the low yield range near the limit of detection. Indeed, we removed 47 (38%) samples which did not pass our relatively lenient filtering criteria. Therefore, the biological signal captured is most likely limited by the sampling methodology.

The natural heritage collections consisting of touched and untouched mollusc and fossil objects allowed us to derive a microbial human touch signature. Increase in this signature was detected on areas of cultural heritage objects with greater exposure to visitors, confirming its validity. Of note, the touch signature demonstrated strong robustness across heterogenous objects, materials, locations, as well as multiple sampling and sequencing rounds.

On an architectural monument like the Ishtar gate in the PM, these results suggest that the microbiome composition reflects the exposure to human interaction. We cannot be certain whether, and if, how excessively, visitors actually touch the monument surface, but a greater level of exposure in lower compared to higher tile levels is obvious. However, even at a smaller physical scale, differentially exposed areas carry distinct microbial profiles as demonstrated at the Sam’al gate lions.

These results suggest that microbial fingerprints can be used to measure, monitor and eventually control exposure to human contact on museum objects.

Interestingly, both the mollusc and fossil objects lost their original microbial fingerprint upon touch. A similar observation was made with the Ishtar gate tiles exposed to visitors that gained the personal microbial fingerprint of subject B upon touch. These observations suggest that upon touch the previous interaction history of the object may get partially overwritten or even lost. In summary, our findings may have implications relevant to heritage provenance or forensic and criminological studies.

## Materials and methods

### Sample Collection

A series of objects exhibited or stored at three cultural heritage institutions in Berlin, Germany (Museum für Naturkunde, MfN; Pergamonmuseum, PM; Rathgen-Forschungslabor, RF) were sampled. The objects were selected to reflect different degrees of exposure to human contact (Table 1). Sampling was carried out with certified DNA-free and sterile nylon-flocked swabs (Copan, 552 C, Brescia Italy), using a minimally invasive methodology previously optimized by Deutsche Sammlung von Mikroorganismen und Zellkulturen (DMSZ) researchers (Flocco et al, in preparation). Briefly, for the first sampling round, 2 by 2 cm areas were swabbed for approximately 90 seconds. During the second sampling round at the PM areas of 0.3m^2^ were swabbed for approximately 3 minutes. After each sampling, swabs were immediately placed in vials and transported to the DSMZ laboratories in Braunschweig and stored frozen at -20 °C till further processing. The same procedure was applied to swabs without sample which were used as negative controls for materials, reagents and processes.

### DNA Extraction, 16S rRNA gene amplification and sequencing

A workflow ‘from swabs to sequence’ for biomolecular analyses of cultural heritage objects, previously optimized by DSMZ researchers (Flocco et al. in preparation), was employed. The main steps and reagents are briefly described here. For the extraction of nucleic acids, swab samples were brought out of the freezer and allowed to thaw at room temperature. DNA was extracted using the QIAamp DNA Micro Kit, (Qiagen) selecting the protocol for isolation of genomic DNA from tissues and proceeding according to manufacturer’s instructions. DNA was quantified using the Quant-iTTM Picogreen dsDNA reagent and kit (Invitrogen) according to the manufacturer’s instructions. The 2-step PCR amplification targeted the V3–V4 region of the 16S rRNA gene with specific primers which also contained an adaptor and barcode sequence. PCR products were purified using the Nucleospin® Gel and PCR clean up kit (Machery-Nagel), according to manufacturer instructions, followed by quantification using the QubitTM dsDNA high sensitivity assay kit (Invitrogen) and pooled equimolarly. The 16S amplicon library pool was cleaned and concentrated using the AMPure XP magnetic beads (Beckman Coulter) and re-quantified using the mentioned Qubit assay. Sequencing was performed using the MiSeq sequencing technology (Illumina Inc., LA Jolla, CA) with 500 cycles generating paired-end sequences with 300 base pairs per read.

### Bioinformatic processing and quality control

Next generation sequencing reads were processed using the Qiime2 software (Bolyen et al. 2019). A Qiime2 object combining reads from all samples from both sampling rounds was created using the *qiime tools import* command. Next, paired-end reads were joined using the *qiime vsearch join-pairs* command (Rognes et al. 2016). Reads were filtered based on quality scores and the presence of ambiguous base calls using the *qiime quality-filter q-score* command (Bokulich et al. 2013). Data was subsequently denoised using the *qiime deblur denoise-16S* command (Amir et al. 2017). Taxonomic classification was performed using the SILVA database (release 132) (Quast et al. 2013) and the *qiime feature-classifier classify-sklearn* command.

### Statistical analysis

Statistical analysis was performed using the R programming language making use of the qiime2R and phyloseq (McMurdie and Holmes 2013) packages. Qiime2 data was loaded into R using the *read_qza* function of the qiime2R package.

Samples with less than 500 total reads or less than 50 unique ASV features detected as well as negative controls were removed from subsequent analysis. The final Qiime2 object after quality control filtering contained a total of 3,184,281 reads across 5,787 unique ASV features and 79 samples. Data was normalized by calculating the square-root of the proportion of total reads assigned to each ASV feature.

We applied independent dimension reduction for various subsets of samples using principal component analysis. For each principal component analysis, ASV features were subset to those detected in at least two samples. Principal component analysis was performed using the *prcomp* function and the scale parameter set to “TRUE”.

Differential abundance analysis was performed using multivariate regression as implemented in the *lm* function. The normalized ASV abundance was used as the dependent variable of the model, while the independent variables were derived from the metadata. P-values were corrected for multiple testing using the *p.adjust* function with the method parameter set to “BH”.

The touch signature was calculated as the proportion of reads derived from the 24 ASV features that were significantly positively associated with touch status in the mollusc and fossil objects.

Data visualizations were generated using the ggplot2 (Wickham 2016), pheatmap and gridExtra R packages.

## Author Contributions

CQ, JV, LS and SS obtained funding for the project. CF and JO performed sequencing experiments. CM, DH, LR, and LS performed bioinformatic processing and data analysis. LS and SS wrote the manuscript.

## Data Availability Statement

All data and analysis code is available at: https://github.com/lkmklsmn/lounsbery.

## Supplemental figures

**Figure S1.**
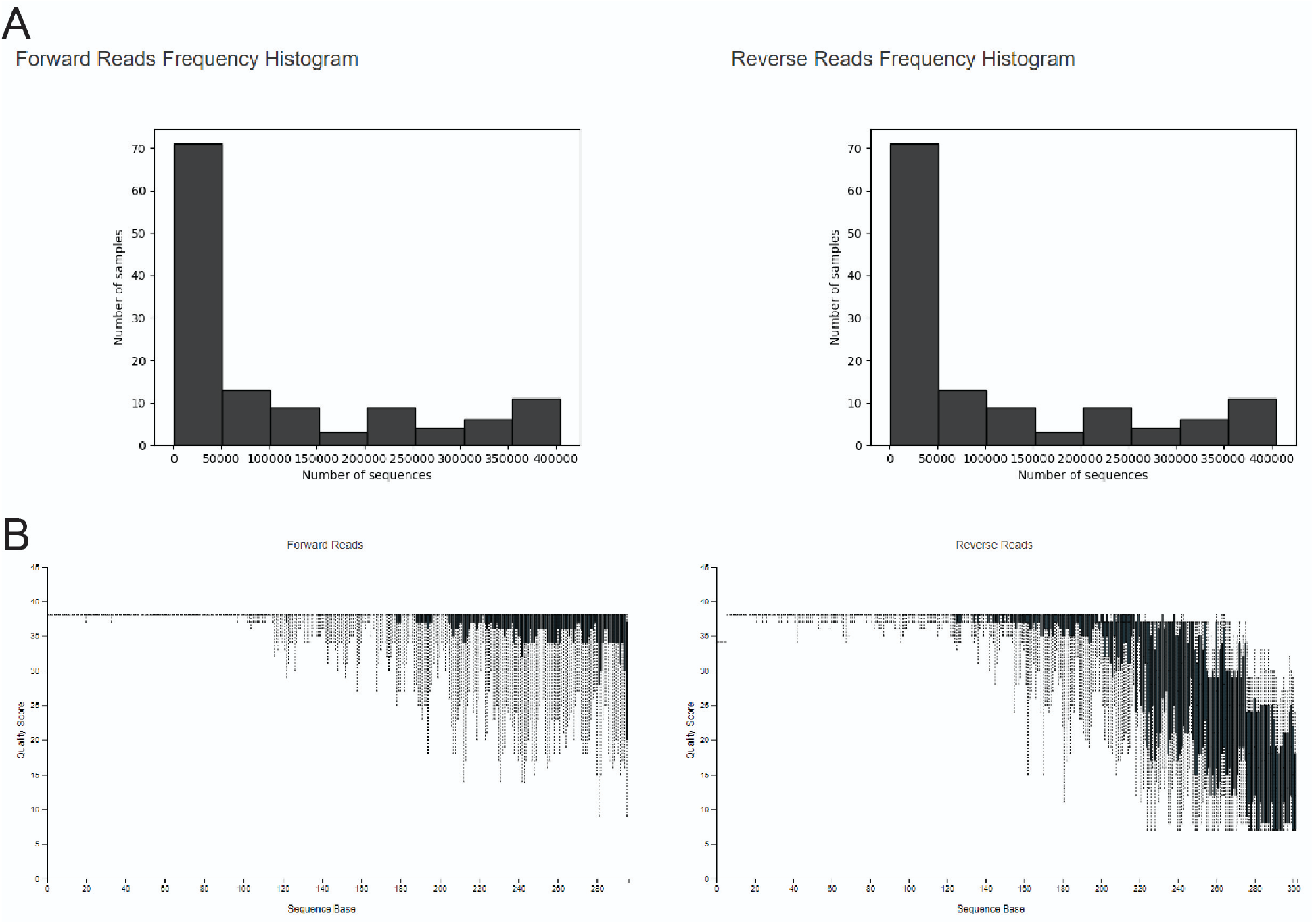
Quality control statistics as output by the Qiime2 software illustrate histograms of demultiplexed sequence counts (A) and quality scores (y-axis) by sequence base (x-axis) (B).

**Figure S2.**
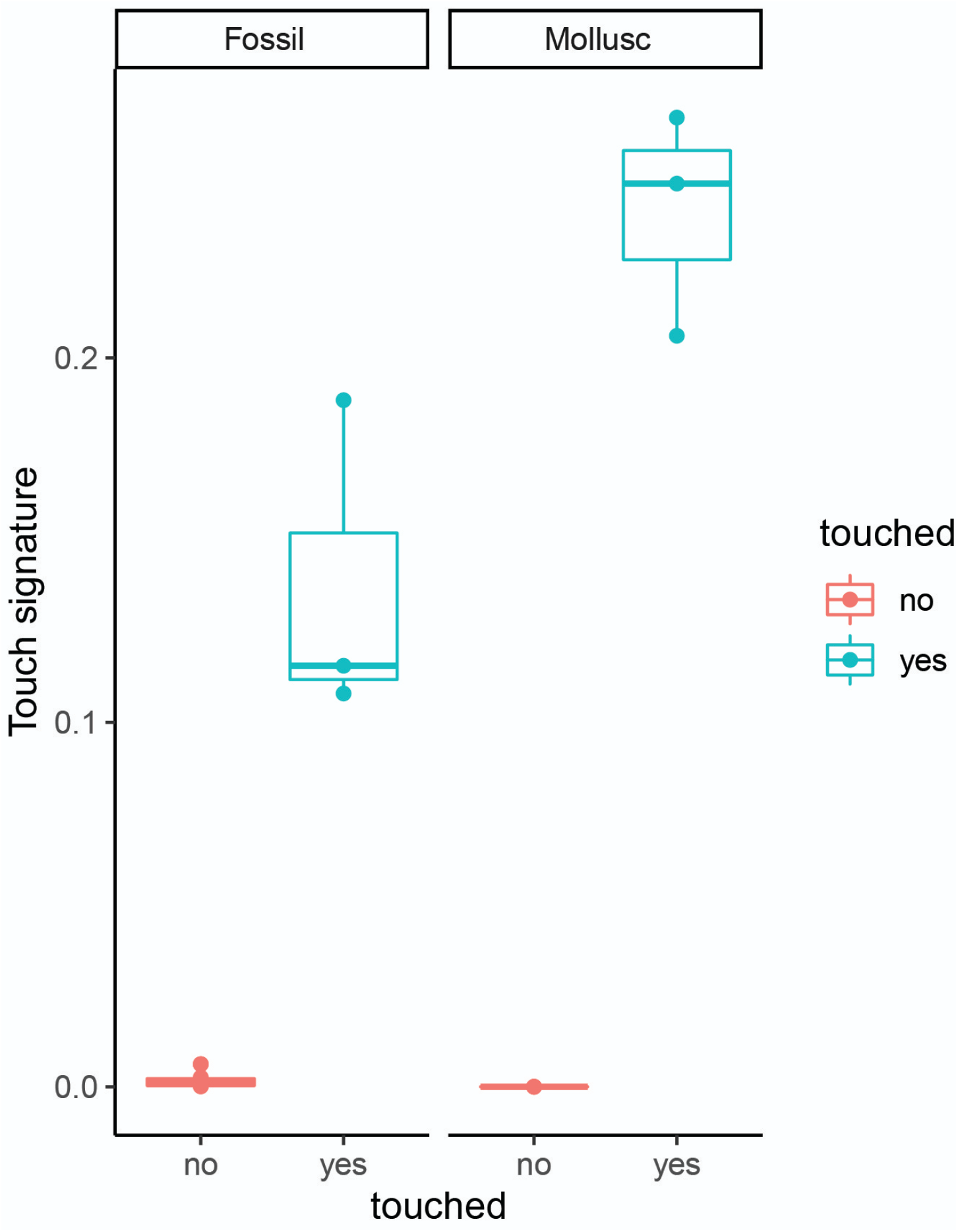
Boxplot displays the levels of the touch signature score across touched (blue) and untouched (red) samples for both molluscs (right) and fossils (left).

**Figure S3.**
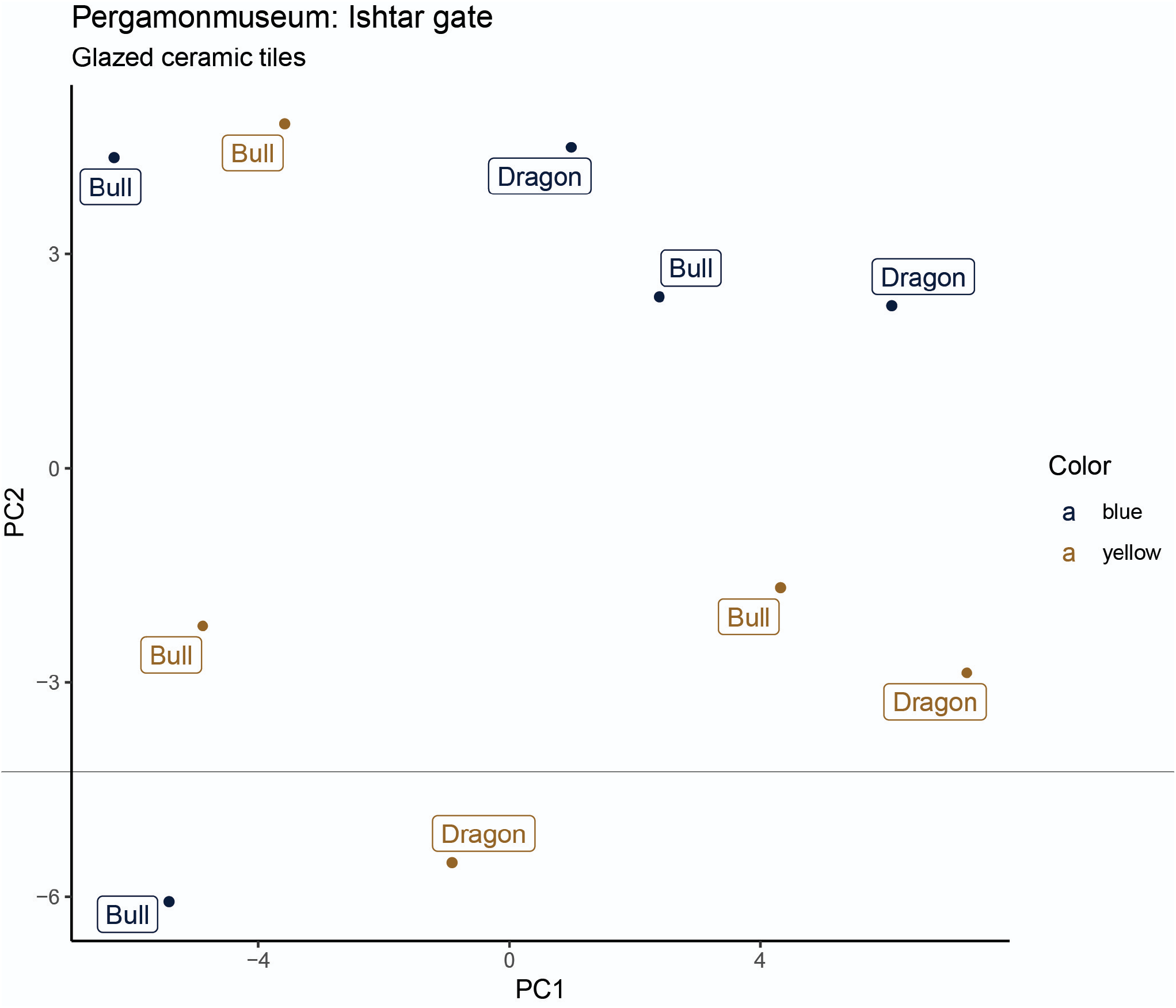
Principal components one (x-axis) and two (y-axis) stratify glazed ceramic tiles from Ishar gate. Color represents the color of the tiles. Illustration figures are labeled.

**Figure S4.**
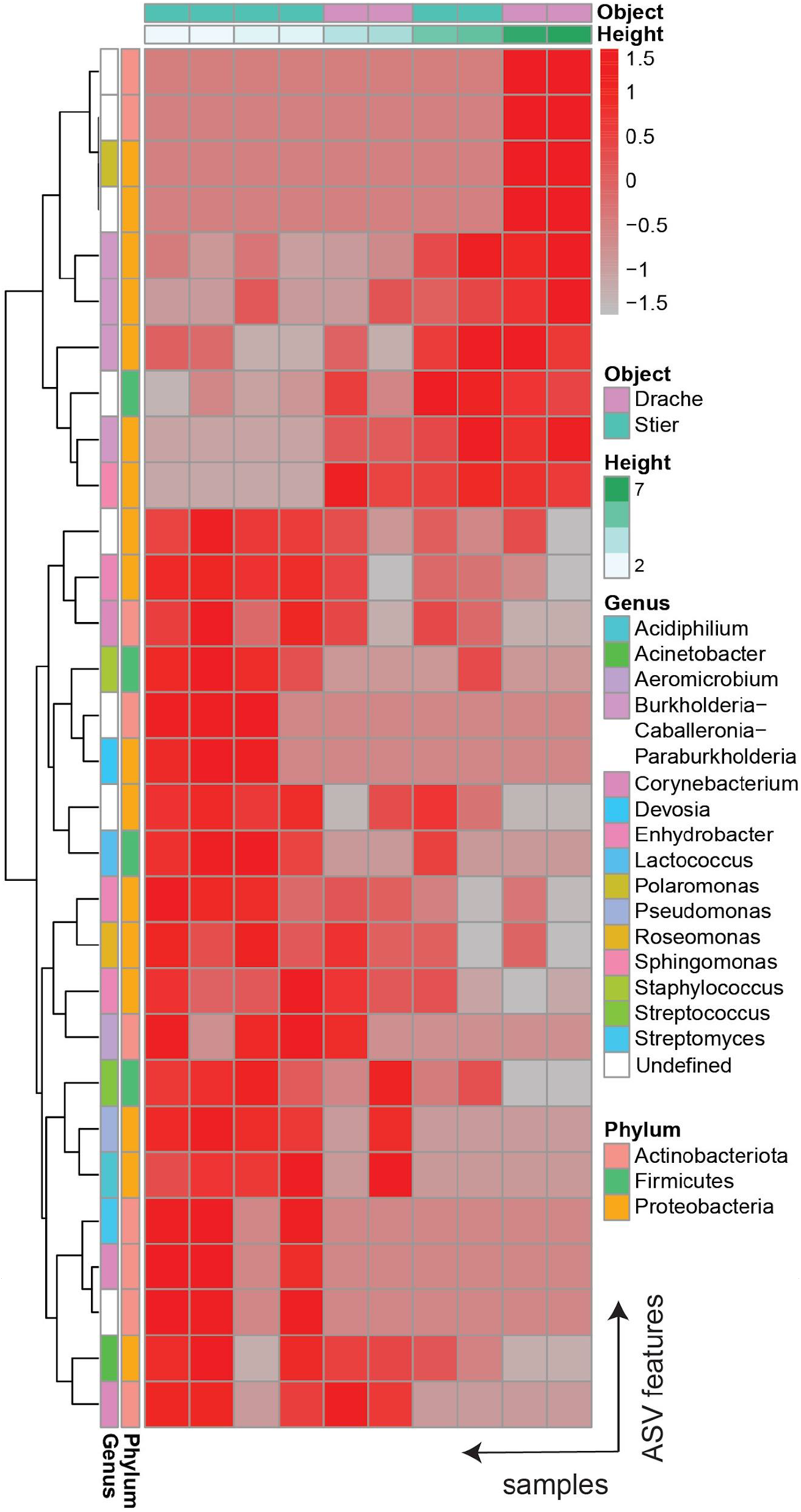
Heatmap illustrates the abundance of top 30 ASV features (rows) associated with height across samples taken from the Ishar gate (columns).

